# Spin coated chitin films for biosensors and its analysis are depended on chitin-surface interactions

**DOI:** 10.1101/215566

**Authors:** Marco G. Casteleijn, Dominique Richardson, Petteri Parkkila, Niko Granqvist, Arto Urtti, Tapani Viitala

**Affiliations:** Drug Research Program, Division of Pharmaceutical Biosciences, Faculty of Pharmacy, University of Helsinki, Finland; BioNavis Ltd., Tampere, Finland; School of Pharmacy, Faculty of Health Sciences, University of Eastern Finland, Kuopio, Finland

**Author notes:** Corresponding author: Marco Casteleijn, Division of Pharmaceutical biosciences and the Drug Research Program, University of Helsinki, Viikinkaari 5 E, 00014 University of Helsinki, FINLAND.

**Keywords:** Chitin, Surface Plasmon Resonance, Chitin Binding Domain, Split-Inteins, Biosensor, Layer Modeling

## Abstract

Chitin, abundant in nature, is a renewable resource with many possible applications in bioengineering. Biosensors, capable of label-free and in-line evaluation, play an important role in the investigation of chitin synthesis, degradation and interaction with other materials. This work presents a comparative study of the usefulness of a chitin surface preparation, either on gold (Au) or on polystyrene (PS). In both cases the most common method to dissolve chitin was used, followed by a simple spin-coating procedure. Multi-parametric surface plasmon resonance (MP-SPR), modeling of the optical properties of the chitin layers, scanning electron microscopy, and contact angle goniometry were used to confirm: the thickness of the layers in air and buffer, the refractive indices of the chitin layers in air and buffer, the hydrophobicity, the binding properties of the chitin binding domain (CBD) of *Bacillus circulans*, and the split-intein capture process. Binding of the CBD differed between chitin on Au versus chitin on PS in terms of binding strength and binding specificity due to a less homogenous structured chitin-surface on Au in comparison to chitin on PS, despite a similar thickness of both chitin layers in air and after running buffer over the surfaces. The use of the simple method to reproduce chitin films on a thin polystyrene layer to study chitin as a biosensor and for chitin binding studies was obvious from the SPR studies and the binding studies of CBD as moiety of chitinases or as protein fusion partner. In conclusion, stable chitin layers for SPR studies can be made from chitin in a solution of dimethylacetamide (DMA) and lithium chloride (LiCl) followed by spin-coating if the gold surface is protected with PS.

## Introduction

Chitin, along with its derivatives, is an increasingly popular biomaterial due to several useful properties, including biodegradability, low immunogenicity, non-toxicity, and biocompatibility [13]. Chitin is the main constituent of the exoskeletons of Arthropods, and is also found in fungal cell walls [3], cocoons of moths [4], diatoms, coralline algae, Molluscae, Protists, Polychaetes, within fish, [5] and marine and freshwater sponges [6, 7], and it is the second most abundant biopolymer after cellulose [3]. Chitin easily be processed in to a number of derivatives, *e.g.* chitosan; these have shown promising applications in a broad range of areas, including food science, medicine, agriculture [8], and the biomedical field [9].

Chitin is an insoluble linear polymer of aminoglucopyrans with extended linear chains of ß-1,4-linked N-acetylglucosamine residues (figure 1), and three polymorphs of this polysaccharide are known [10, 11]. It exists in three structural forms: α-, ß- and γ-chitin that have different mechanical properties depending on the arrangement of the polymer chains. The α-form represents an alternating antiparallel arrangement of polysaccharide chains (e.g. crustaceans [12]), the ß-form is a parallel chain arrangement (found e.g. in squid), and in the γ-form two parallel chains statistically alternate with an antiparallel chain (found e.g. in fungi) [4, 13]. The α-chitin isomorph is most abundant and formed by: enzymatic polymerization [14], *in-vitro* biosynthesis [15, 16], and recrystallization from solution [17]. Chitin is more than 50% acetylated, while chitosan is primarily deacetylated [11]. Shrimp chitin has been reported to be > 95% acetylated [18].

**Figure 1.**
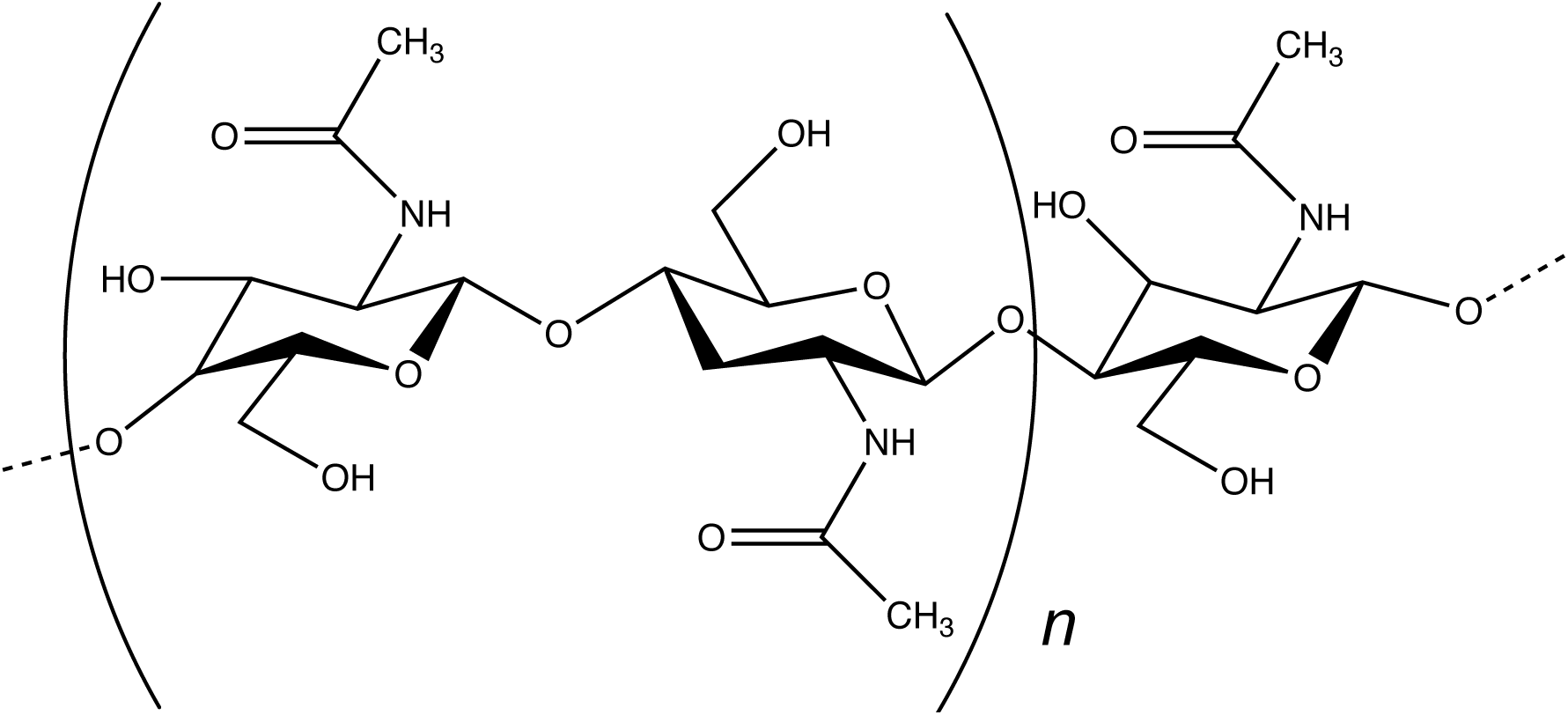
Chemical structure of chitin. Chitin polymers derived from shrimp shell have a typical degree of polymerization of 493 – 605 [64].

The use of chitin, and to a greater extent, chitosan films as constituents of biosensors have gained interest as the direct study of chitin is important in gaining insight into enzyme degradation pathways, interactions with natural composite materials (e.g., proteins and polysaccharides), and mechanisms behind biocompatibility and biofunctionality [19, 20]. Chitin and chitosan are useful transducer surface modifiers and found many applications as a electrochemical (bio-)senor component, for example in Pt disk electrodes for choline-sensing, supercapacitors [21] for nucleic acid analysis (e.g. DNA microarrays), immunosensors (e.g. biomarker analysis), enzyme activity, and detection of trace elements (for comprehensive reviews please read Suginta et al. *2013* [11] and Kim et al. *2015* [22]). In addition, chitin can be easily modified via its hydroxyl and acetylamido groups for future biosensor advancements. Chitin has advantageous mechanical properties, such as a high mechanical stability (tensile strengths between 38 – 146 MPa have been reported) and a high thermal stability (e.g. shrimp shell chitin’s thermal degradation is between 290 – 440 °C), though these properties are underused in industry [5].

In order to create chitin surfaces with representative properties similar to its natural structure, chitin needs to be dissolved first, and then deposited onto the sensor surface. One approach is the use of biomimetic methods, for example to use naturally occurring biocomposites. For instance, instead of spin-coating chitin on a silicon surface [23], silica-chitin based biocomposites, or other biocomposites, could be used and it is available from various sources and can even be obtained in-vitro [24–28]. A newly emerging area is the use of natural sources from environments where life is found under extreme heat or pressure, also referred to as extreme biometrics [5]. The solubility of chitin depends on the pH and ionic strength of the solvent used and is influenced by the percentage and distribution of acetylated (and deacetylated) moieties along the backbone of the polysaccharide [11]. It is high in crystallinity and does not dissolved in hot water [27]. The most common method to solubilize chitin is the use of dimethylacetamide (DMA) and lithium chloride (LiCl), already demonstrated at the beginning of the last century [29, 30]. Recrystallizing chitin from a 5% LiCl in DMA solution by precipitation had an essentially crystalline form as the chitin prior to solution (be it the α- or the ß-form), though the α-form can be slightly disordered [23]. Alternative methods for the solubilization of chitin have been reported, such as using other polar solvents: LiCl/*N*-methyl-2-pyrolidone, chloride dihydrate/methanol [31], and hexofluoroisopropanol (HFIP) [32], or ionic liquids, such as 1-allyl-3-methylimidazolium bromide ([Amim]Br), [C_2_mim][OAc], and 1-butyl-3-methylimidazolium acetate [31]. Thin chitin films are interesting for studies of chitinases which can convert chitin into cheap products such as N-acetylglucosamine for biomedical use [2]. Chitin films on silica surfaces by means of spin-coating were 50 nm thin, but rough [33]. Furthermore, a smooth, thin layer of amorphous chitin was obtained after the spin coating of trimethylsilyl chitin onto a silica or gold surface after which the films were regenerated to amorphous chitin [2].

One analytical method, which is highly compatible with thin layers is surface plasmon resonance (SPR). It is a highly sensitive, non-invasive, and label-free technique that is used extensively for surface interaction studies between polymers and proteins [34]. In addition, it has been used to evaluate a chitin-coated surface in the past using ionic liquids as a solvent [2]. However, no simple protocol for the preparation of thin chitin layers using DMA/LiCl as solvent on SPR gold sensors has been established yet.

In this contribution, we demonstrate that the precipitation of chitin, derived from shrimp shell, using the common DMA/LiCl solvation method onto a gold surface for subsequent SPR analysis poses a challenge due to the surface properties of gold, but can be protected by an additional coating of polystyrene prior to chitin coating. The composition, thickness, available binding surface, surface hydrophobicity, and ability to interact with a CBD of chitin were characterized with multi-parametric surface plasmon resonance (MP-SPR), scanning electron microscopy (SEM), and contact angle goniometer with two different SPR sensors: bare gold and a polystyrene coated gold. The binding constants of a chitin binding domain fused to a split intein and its split intein counterpart were determined for both systems, as well as the differences in behavior of chitin on gold versus polystyrene after spin coating.

## Methodology

### Materials

Unless specifically mentioned all commercial chemicals (dimethylacetamide (DMA), LiCl, hexofluoroisopropanol (HFIP), EDTA, DTT, NaN_3_, triethanolamine hydrochloride (TEA), Tris, sodium chloride, ampicillin, Isopropyl β;-D-1-thiogalactopyranoside (IPTG), magnesium chloride, and chitin polymers derived from shrimp shell (C7170, lot: 051M7013V) were obtained from Sigma-Aldrich (USA) with a reported acetylation degree of 98 ± 12 % [18]. Reaction buffers and media used were composed as follows: Buffer A: 100 mM TEA, 1 mM EDTA, 1mM DTT, 1mM NaN_3_, pH 7.6 (at 25° C). Buffer B: 20 mM TEA, 1 mM EDTA, 1mM DTT, 1mM NaN_3_, pH 8 (at 25° C). Buffer C: 20 mM Tris/HCl, 100 mM sodium choride, pH 7.0 (at 25° C). Buffer D: 50 mM Tris/HCl, 50 mM NaCl, pH 8.0 (at 25°C). DNAse and RNAse were obtained from Epicentre (Madison, WI, USA). Phenylmethylsulfonyl fluoride (PMSF) was obtained from ThermoFisher Scientific, USA. EnPressoB medium was obtained from BioSilta Ltd. (U.K); currently available via EnPresso GmbL (Berlin, DE).

### DNA methods

The eGFP gene from the pEGFP-C1 plasmid (Clontech, USA) was cloned into the pTYB21 plasmid, containing the *Sce*VMA1 gene (NEB, USA) to yield the eGFP-SceVMA1 Intein-CBD construct with use of Gibson Assembly^®^ (NEB, USA) according to manufacturer’s instructions. The eGFP gene was also cloned into a pMMRSF17 plasmid that contained the NpuDnaE Intein gene in the same way (the full construct was a gift from Hideo Iwai based on Addgene plasmid # 20178). In order to generate the CBD-NpuDnaE_NΔ16_ gene, the CBD of the chitinase A1 (CBD_ChiA1_) enzyme of *Bacillus circulans* WL-12 was isolated from the pTYB21 plasmid (NEB, USA) using the following primers in a gradient-PCR reaction using Dynazyme II polymerase (ThermoFisher Scientific, USA): ‘5-GGAATTCCATATGACAAATCCTGGTGTATC-3′ and 5′-GGATCCGAATTCTTGAAGCTGCCACAA-3′ (fragment A) and the NpuDnaE_NΔ16_ gene isolated from the pMMRSF17-NpuDnaE with primers 5′-GAATTCGGATCCGCCTTAAGCTATGAAACGAA-3′ and 5′ -CGTTAAGCTTATTGAAGCTGCCACAAGG-3′ (fragment B). The last primer introduced a mutation (Cys **→** Ala) at position one of the NpuDnaE_NΔ16_ gene and a linker sequence (EFGS) between the CBD and NpuDnaE_NΔ16_ in the final construct in pET21a (see below). In order to generate the NpuDnaE_C16__ eGFP gene, the NpuDnaE_C16_ was isolated from the pMMRSF17-NpuDnaE plasmid using primers 5′-GGAATTCCATATGGGTCATAATTTTGCACTC-3′ and 5′-CCCTTGCTCCCATATTAGAAGCTATGAAG-3′ (fragment C) and the eGFP gene was isolated from pTYB21-eGFP-Intein-CBD plasmid described above with primers 5′-TCATAGCTTCTAATATGGTGAGCAAGGGCGA-3′ and 5′-TTTCAAGCTTTTACTTCTACAGCTCGTCCATGC-3′ (fragment D). In an overlap extension gradient-PCR (42° C – 60° C; 15 cycles) fragments A and B were combined with use of Dynazyme II polymerase (ThermoFisher Scientific, USA). Fragments C and D were combined in similar way (38° C - 45° C; 15 cycles). Faint bands in the latter were amplified with PCR and all reactions were cleaned up with NucleoSpin^®^ Gel and PCR Clean-up columns (Macherey-Nagel, DE). The CBD-NpuDnaE_NA16_ gene and the NpuDnaE_C16__ eGFP gene were then separately cloned into pET21a expression vectors (Novagen, Merck KGaA, USA) after digestion with NdeI and HindIII (ThermoFisher Scientific, USA) and ligation with T4 ligase (ThermoFisher Scientific, USA). The complete DNA sequences of all genes were verified by gene sequencing (GATC, DE). *Escherichia coli* NEB5-alpha competent cells (New England Biolabs, USA) were transformed and used for plasmid propagation [35]. Plasmids were then transformed separately to *E. coli* strain BL21(DE3) (Invitrogen, USA) for protein production.

### Protein expression and purification

Expression of eGFP-*Sce*VMA1 Intein-CBD, CBD-NpuDnaE_NΔ16_ and NpuDnaE_C16_-eGFP was carried out in BL21(DE3) *E. coli* cells. First the transformants were grown o/n in LB Medium with 100 μg/ml ampicillin and 1% (w/v) glucose at 30°C from glycerol stocks; 2 ml o/n culture was used to inoculate 50 ml of EnPressoB medium [36] with 100 *μ*g/ml ampicillin at 30° C; 225 rpm (1” amplitude shaker) according to manufacturer’s instructions in high yield flasks [37]. Cells were induced with 0.4 mM IPTG after 24 h and grown for another 24 h under the same conditions. Cells were separated from the media via centrifugation (16 000 * g for 15 min) and then lysed by freezing o/n at −20° C in the presence of DNAse (2.5 U/ml), RNAse (2.5 U/ml), Lysozyme (15 μg/ml), 0.1

mM PMSF (ThermoFisher Scientific, USA), and MgCl_2_ in buffer D. The insoluble fraction was removed by centrifugation (15 min at 9000 * g). The soluble protein fraction of CBD-NpuDnaE_NΔ16_ was loaded onto a 1 ml HiTrap DEAE FF column (GE healthcare USA) and eluted with buffer D at incremental sodium chloride concentrations (100 mM, 150 mM, 200 mM, 250 mM, 500 mM). The soluble pooled and concentrated protein fractions of NpuDnaE_C16_-eGFP were heated at 70° C for 20 min, insoluble proteins removed by centrifugation (15 min; 16 000 *g), and the supernatant loaded onto a size exclusion chromatography (SEC) HiLoad Superdex 200 prep grade filtration column (GE healthcare, USA) using buffer A at 0.5 ml/min and protein fractions collected. The purity of all fractions was evaluated by SDS-PAGE analysis (AnyKd gels, BioRad, USA) and fractions of the target protein size were pooled. Eluate buffer A was replaced by buffer C using a 5kD NMWL membrane at 2800 rpm (Merck-Millipore, USA). Protein concentrations were determined by OD_280_; Molecular weight and the ε was calculated with the ProtParam tool [38]. Hereafter we will abbreviate CBD-NpuDnaE_NΔ16_ as CBD-IN_N_ and NpuDnaE_C16_-eGFP as IN_C_-eGFP.

### Chitin coating

Chitin was dissolved in DMA and 5% LiCl according to Austin P.R. (1984) [39] at a final concentration of 0.1% (w/v). Bare gold sensors were spin-coated with 100 μ! for 60 sec. at 3600 rpm; polystyrene coated senors (BioNavis Ltd., Ylöjärvi, Finland) were spin-coated by adding 100 μl to the sensor and then after 60 sec. the sensor was spin-coated at 60 sec. at 3600 rpm. The sensors for SPR were used only once and spin-coated immediately before use, since LiCl leached the gold off of the bare sensors after drying for longer periods at 60° C, and subsequent rehydrating (fig. 1).

For scanning electron microscopy imaging the sensor was spin-coated prior to placement in the vacuum chamber of the SEM microscope (application chamber) to determine the structure before applying buffer. Sensors were then placed within 10 minutes in a SPR Navi™ 200 (BioNavis Ltd., Ylöjärvi, Finland) instrument and buffer was applied to the flow cell area at 20 °C, with a flow rate of 20 μL/min. Samples were allowed to air-dry for at least 15 minutes before SEM imaging of the flow cell area and the areas outside the flow cell under the same conditions.

### MP-SPR measurements

Measurements were performed with a multi-parameter SPR Navi™ 200 (BioNavis Ltd., Ylöjärvi, Finland) instrument. The setup was equipped with two incident laser wavelengths, 670 nm and 785 nm, two independent flow channels, inlet tubing and outlet (waste) tubing. Both of the flow channels were measured in parallel with 670 nm and 785 nm incident light. The measurement temperature was kept constant at 20 °C. The chitin coated SPR sensors were placed in the flow-cell and SPR spectra were recorded in air after which buffer B was introduced into the flow-cell; the flow rates used for the CBD-IN_N_ interaction with chitin and for IN_C_-eGFP interaction with CBD-IN_N_ were 20 μL/min. Chitin layer data in air was fitted in LayerSolver 1.2.1 (BioNavis, FIN) with default parameters and simulation mode 0 for the Au layer, and mode 2 for all other layers. The buffer (due to the presence of EDTA) prevented damage to the bare-gold sensors during the SPR measurements due to removal of LiCl. The chitin coated gold and polystyrene sensor slides were first subjected to buffer B for 10 – 30 min until a stable baseline was achieved. In the second phase, CBD-IN_N_ was injected into both flow channels for 8 minutes at increasing concentrations (490, 981, 1961, 3922, 7854 nM, equivalent to 10, 20, 40, 80, 160 ug/ml) to determine the binding constants for both systems. Based on these results, CBD-IN_N_ was bound to the new chitin coated sensors at 0. 5 and 5 *μ*g/ml of protein during a 5-minute injection. IN_C_-eGFP protein was then injected in both channels for 5 minutes at increasing concentrations (17.5, 175, 350, 701, 1401, 2803 nM, equivalent to 0.5, 5.0, 10.0, 20.0, 40.0, and 80.0 *μ*g/ml) to determine the binding constant of the split-intein pair for both SPR sensors. The data was fitted in TraceDrawer software v1.6 (BioNavis, FIN & Ridgeview Instruments, SWE) according to the OneToTwo model to obtain the association rates, dissociation rates, and the binding constants after evaluations of other relevant models.

### Scanning Electron Microscopy

SEM images were recorded using an FEI Quanta 250 Field Emission Gun Scanning Electron Microscope under high vacuum (< 6 × 10^-4^ Pa) to prevent deterioration of the chitin layer due to moisture in the air.

### Contact angle measurements

Au, PS, Au-chitin, and Au-PS-chitin surfaces were prepared as described above. The Au sensor surface was cleaned in two ways: (1) plasma treated as above and (2) boiled for 15 min in and NH_3_ (30%)/H_2_o_2_ (30%)/H_2_o (1:1:5, v/v) oxidizing solution. A syringe and needle were washed with 70% ethanol and rinsed with fully deionized water (Milli-Q water) and the camera was adjusted before each measurement. The shape of the water droplet on the surface was recorded with a CAM 200 optical contact angle meter (KSV instrument LTD) equipped with CCD video camera module and the drop volume was constant during the whole measurement (static mode). The drop height prior to the measurement was 200 pixels (approx. 8 *μ*l of Milli-Q water), and recording started automatically upon detection of the drop on the surface with a frame interval of 1 second for 60 frames. Contact angels were calculated according to the Young-Laplace fitting model using the Attension theta 4.1.0 software (KSV instruments LTD). After the contact angle was stable (i.e. in the case of Au and PS) with out chitin) the mean of frames from 45 second to 60 second was calculated. For the contact angle of water on the chitin surfaces (dry or wet) the rate of the contact angle change was different, but lower than 1° at the end of each measurement.

## Result and Discussion

### Surface properties of the spin coated sensors

We observed in PS 96-well plates (Corning, USA) with (a) 0.1% chitin solution in DMA/5% LiCl and (b) 0.1% chitin solution in HFIP [32], and subsequently dried overnight in an oven with a fan at 60° C, that (i) chitin precipitated out of DMA/5% LiCl formed large crystal structures and that non-reproducible amounts remaining in the wells after washing with vigorously with water, (ii) chitin precipitated out of HFIP formed a layered gel, which after upon adding water detached from the PS surface, (iii) eGFP-intein-CBD (IMPACT™ kit [40], New England Biolabs, USA) was able to bind and release eGFP according to the kit instructions from crude cell lysates (data not shown). We also used the eGFP-intein-CBD crude lysate to determine that the CBD does not bind to Au (data not shown). Therefore, in order to study chitin binding and the binding/release of split-inteins, SPR Au sensors coated with chitin were initially dried overnight at 60° C. However, the gold was leached from the retrieved SPR sensors after the experiment (i.e. the gold and chitin layer detached from the glass surface; figure 2). For this reason, all binding experiments here reported were performed with buffer B in order to scavenge the LiCl from the surface. In addition, PS coated Au SPR sensors were utilized as well and directly compared. An alternative method, as presented by Kittle et al. *2012* [2] for preparing chitin-SPR sensors produces excellent uniform chitin layer, however the methods we propose requires less experimental steps, and therefore simplifies the use of chitin as a substrate for SPR sensors.

**Figure 2.**
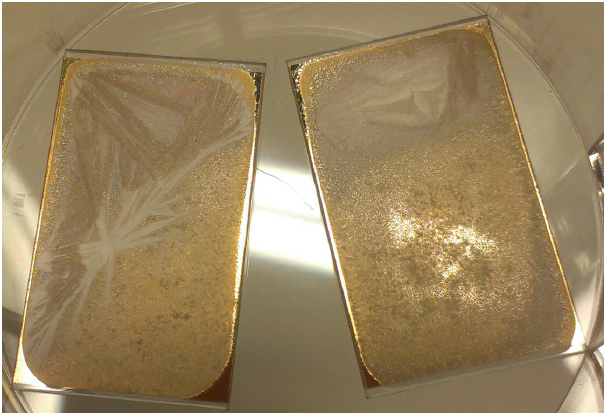
Au leaching of SPR chips due to LiCl in the 0.1% chitin DMA/5% LiCl solution. White areas are void of gold.

The cleaning protocol of gold has an effect on the hydrophobicity of the surface (table 1). Au surfaces boiled for 15 min in a NH_3_ (30%)/H_2_o_2_ (30%)/H_2_o (1:1:5, v/v) oxidizing solution were more hydrophobic than plasma cleaned surfaces. Since it had been shown that: (i) more oxidized gold surfaces are more hydrophilic [41], and (ii) the intention of spin-coated chitin is to interact with its surface more effectively by sampling more hydrogen bonds outside its internal structure [42, 43] all surfaces were plasma treated prior to spin coating. The polystyrene surface can be easily oxidized through plasma treatment [44].

**Table 1.**
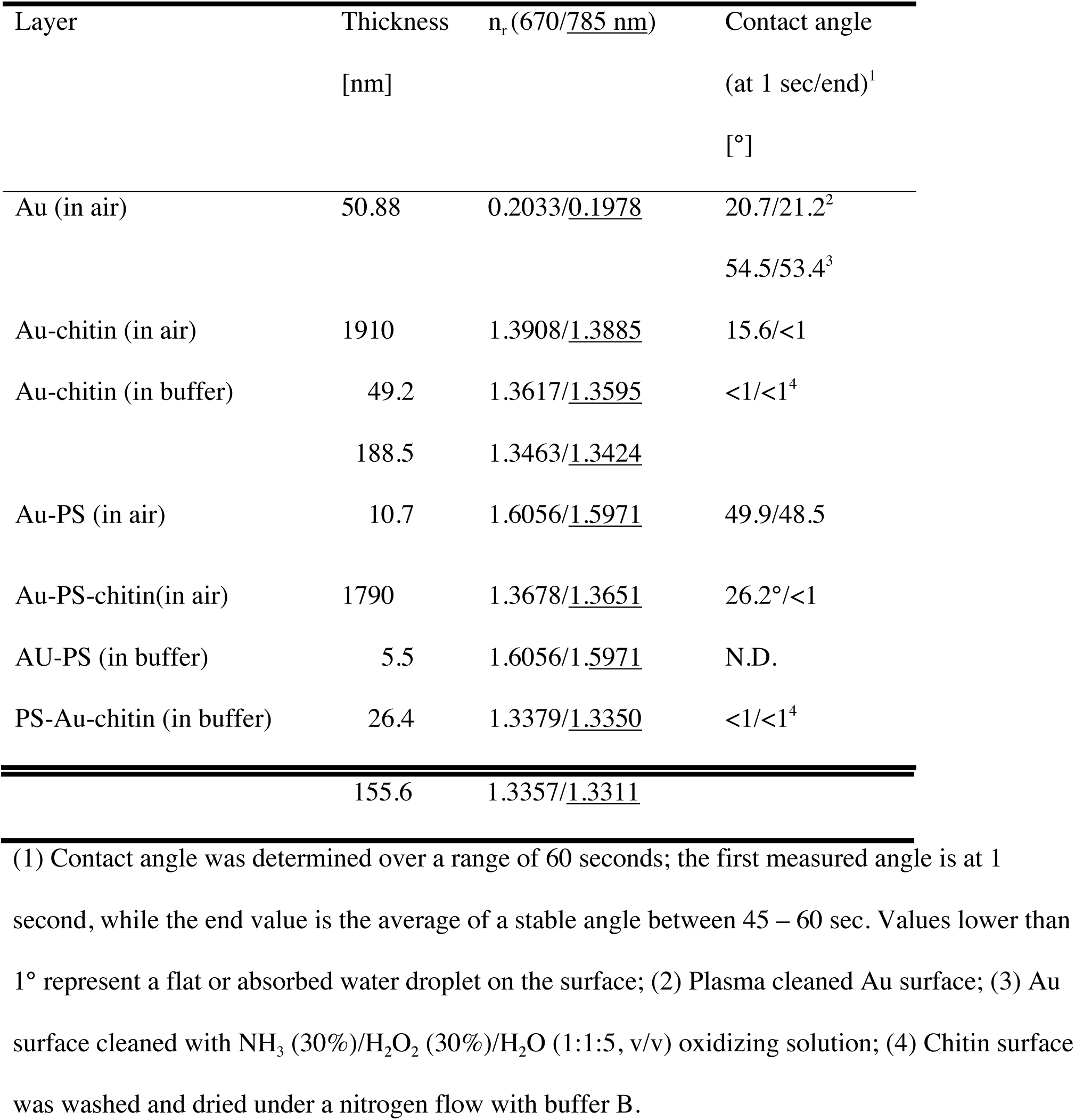
Surface properties of the SPR chips, before and after coating.

The hydrophobicity of the surfaces was evaluated with a goniometer to determine the contact angle of a water droplet during the first 60 seconds of contact. The chitin surfaces were either spin-coated and then dried under a nitrogen flow or spin-coated, wetted with a few drops of buffer B and then dried under a nitrogen flow (Chitin in air or Chitin in buffer in table 1 respectively). On the Au and PS surfaces the water droplet formed was stable throughout the measurement (figure 3), while the water droplet on pre-wetted chitin surfaces immediately spread over the surface. On dried chitin surfaces the water droplet behaved differently than on the Au-chitin surface or the PS-chitin surface. In Au-chitin the contact angle decreased under 10° after 2 seconds and reached a value below 1° within 30 seconds, while the contact angle of a water droplet on PS-chitin was initially higher, the droplet spread slower, and a thicker water layer remained (figure 3).

**Figure 3.**
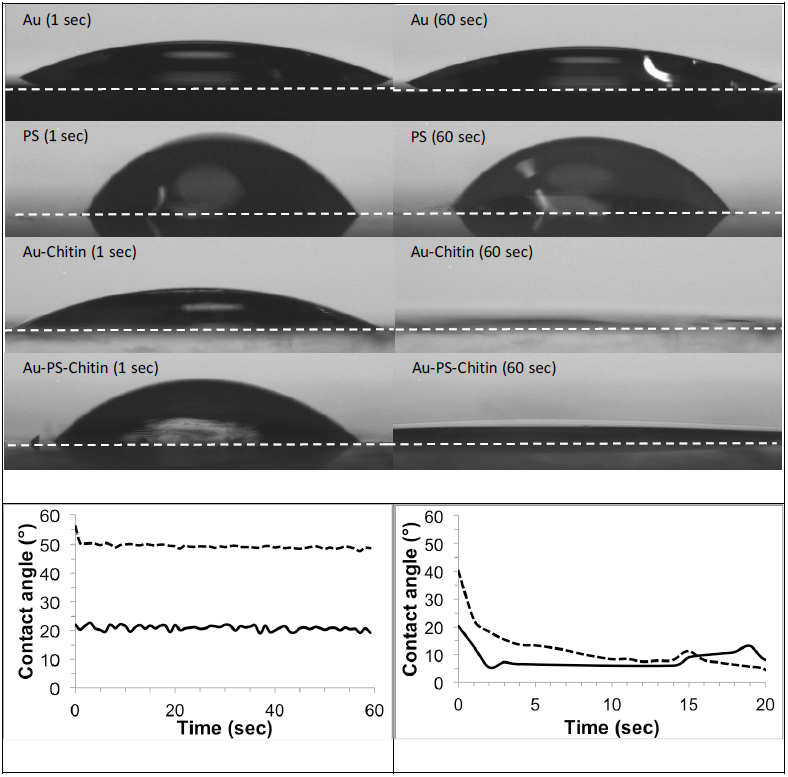
Water droplet behavior during contact angel measurements. The white dashed line in the upper panels depicts the droplet-surface interface. Droplets on Au (solid line) or Au-PS (dashed line) were stabile (bottom left), the droplet on dry Au-Chitin surface (solid line) reached a contact angle below 10° after 2 seconds, while the droplet on the dry Au-PS-Chitin surface (dashed line) reached a contact angle below 10° after 20 seconds (bottom right). All surfaces depicted here were plasma cleaned prior to measurements or spin-coating.

The thickness of the chitin layers directly after spin-coating was determined with SPR in air (table 1 & figure 4) by using the Layersolver software based on Granqvist *et al.* [45]. First the Au layer was measured and modelled (figure 4A) after which chitin was spin-coated on the surface and the thickness determined. Due to the high thickness waveguide-fitting for one wavelength would suffice, but the measured SPR spectra at both wavelengths fit the models well. Therefore, we always fitted both wavelengths in one model to enhance accuracy. The PS coated Au sensor was determined in similar fashion: first the thickness of PS was determined, after which similar model parameters of the chitin on Au-model were applied to obtain the Au-PS models in air. From the models it is clear that the layer’s refractive indices (*n*) in air are much lower than reported for shrimp chitin, which is reported at *n* = 1.61 [46]. This behaviour has previously been observed for spin-coated chitin for which the refractive index can vary greatly between *n* = 1.30 - 1.47 depending on the solvent used [47]. In our case, buffer B at pH 8.0, needed for the protein stability to study their interactions with chitin, may have had an influence on the refractive index of spin-coated chitin.

**Figure 4.**
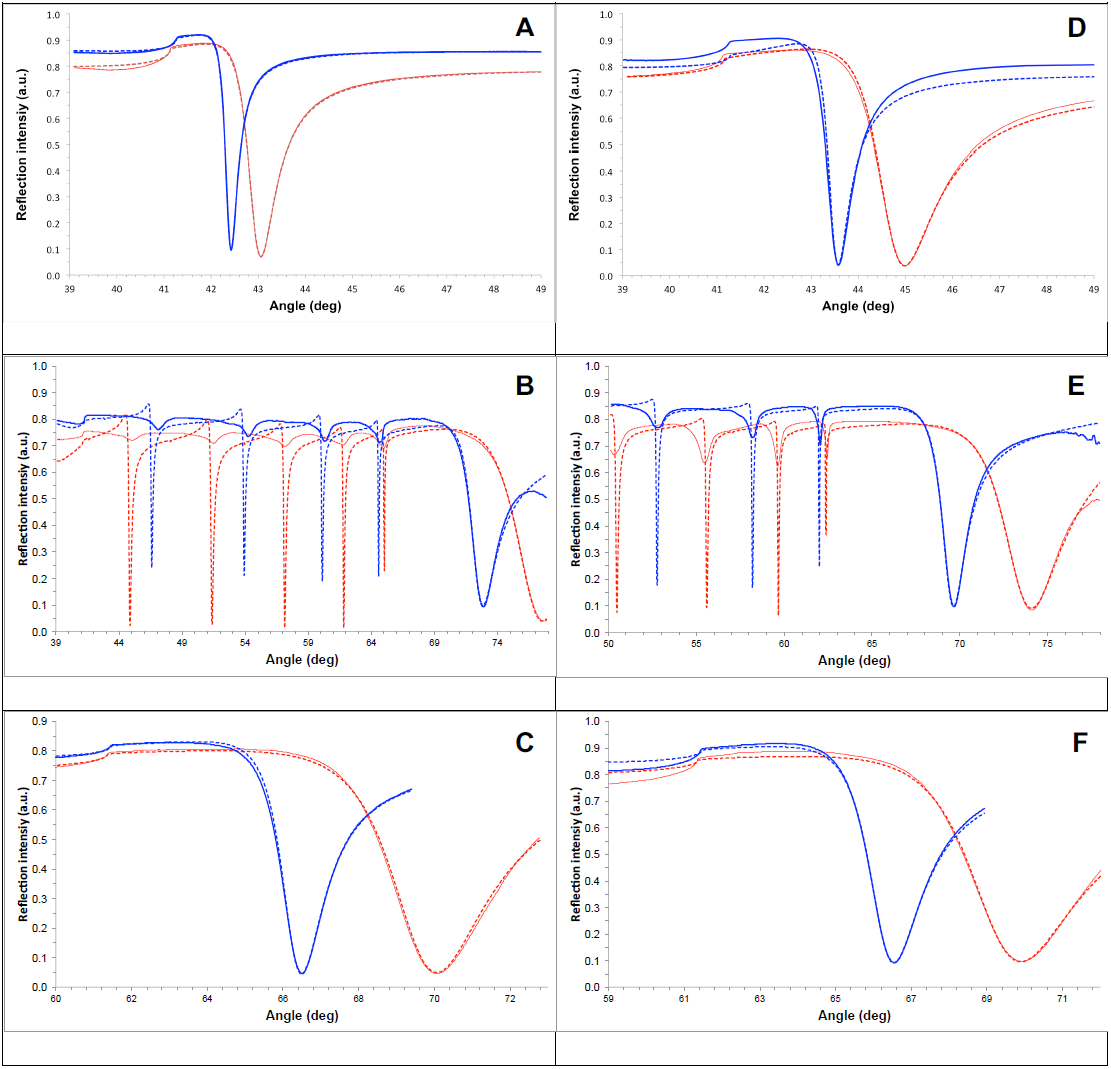
SPR full angle scans of (A) pure SPR sensor background, (B) spin-coated chitin on Au, (D) PS on Au, and (E) spin-coated chitin on PS on Au; 670 nm: red solid line, 785 nm: blue solid line with corresponding Layer solver fits (dashed lines). The optical parameters used for the Layer solver fitting are listed in table 4.

**Table 4.**
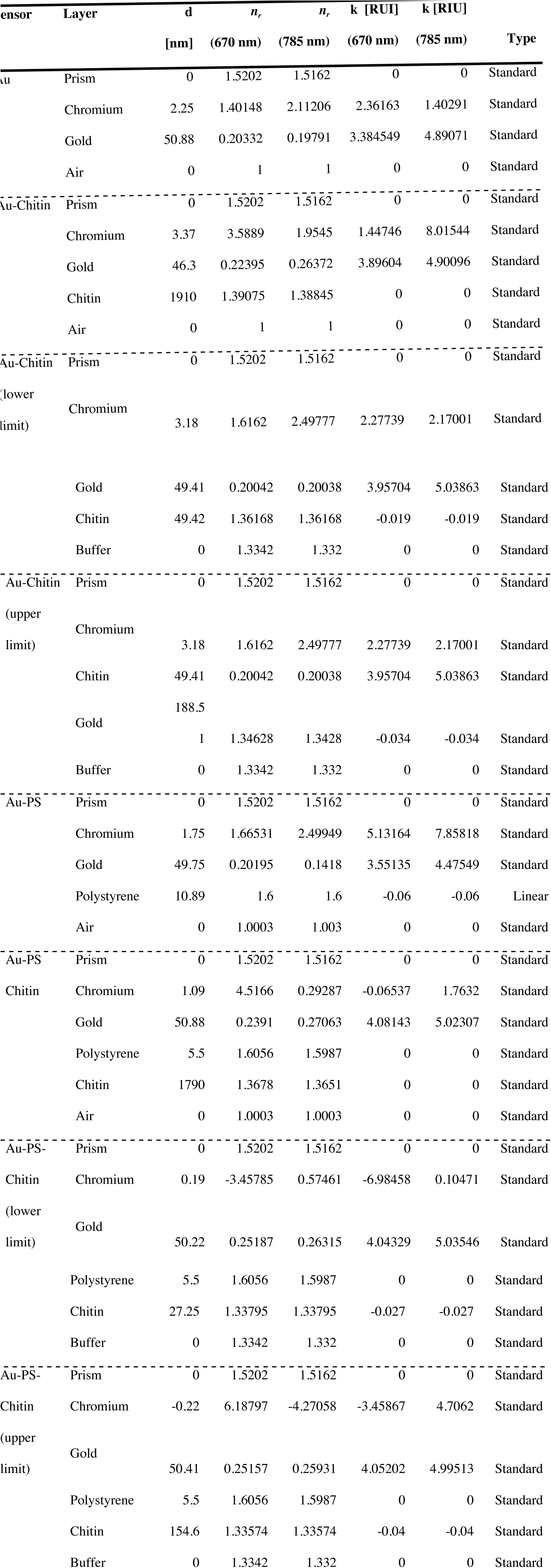
Optical parameters for the layers used in the Layersolver software for figure 4.

In order to determine the refractive index of chitin after the introduction of buffer, the total internal reflection (TIR) angle was manually fitted in the Layersolver software for the Au-chitin layer. Based on the principle that the bulk effect in the SPR curves is constant, while we assume the Au layer to be constant, the changes in the TIR angle-shape during the stabilization of the TIR angles and overall SPR signal are indicative to the changes in thickness. In addition, it is known that hydrogels, such as polyacrylamide gels have a lower refractive index than the starting material [48]. We observed a hydrated state after washing the spin-coated chitin surface with buffer B prior to the contact angle measurements (fig. 3). Overall, the refractive index of chitin can differ greatly depending on how the polymers are precipitated [47]. We disagree with the assumption of Kittle *et al.* [2] that the refractive index of spin-coated chitin is 1.51 and thus we used the change of the shape of the TIR angle to fit the theoretical maximum thickness (fig. 5) and a resulting n-value for both wavelengths for the Au-chitin layer. Thereafter, we fitted the minimal thickness and refractive index for the experimentally SPR curves manually with the Layersolver software. The results are a realistic range of the thickness and refractive indices for both wavelengths. Since we did not observe the phenomena of the TIR angle-shape changing after the introduction of buffer B over time regarding the Au-PS-chitin surface, we took the minimum and maximum refractive indices determined with the Au-chitin models in buffer, and used them as initial values to manually fit the experimental data as above to find a realistic range of the thickness and *n*-values for the Au-PS-chitin layer in buffer. All four resulting models fit the experimental data the best (fig. 4C and 4F). In addition, an apparent change in the thickness of the PS layer had to be corrected for in the models. Therefore we fitted the sensor parameters simultaneously with the layer (table 1). It is known that DMA damages PS [49], however as we observed in Scanning Electron Microscopy (SEM) imaging (fig. 8C and 8D), enough PS remained during the short period of interaction with DMA in the spin-coating process. Since DMA is removed during spin-drying [50] and the PS is not severely damaged during the spin-coating process, the deposited LiCl on the surface is not interacting with Au as observed in figure 8C.

**Figure 5.**
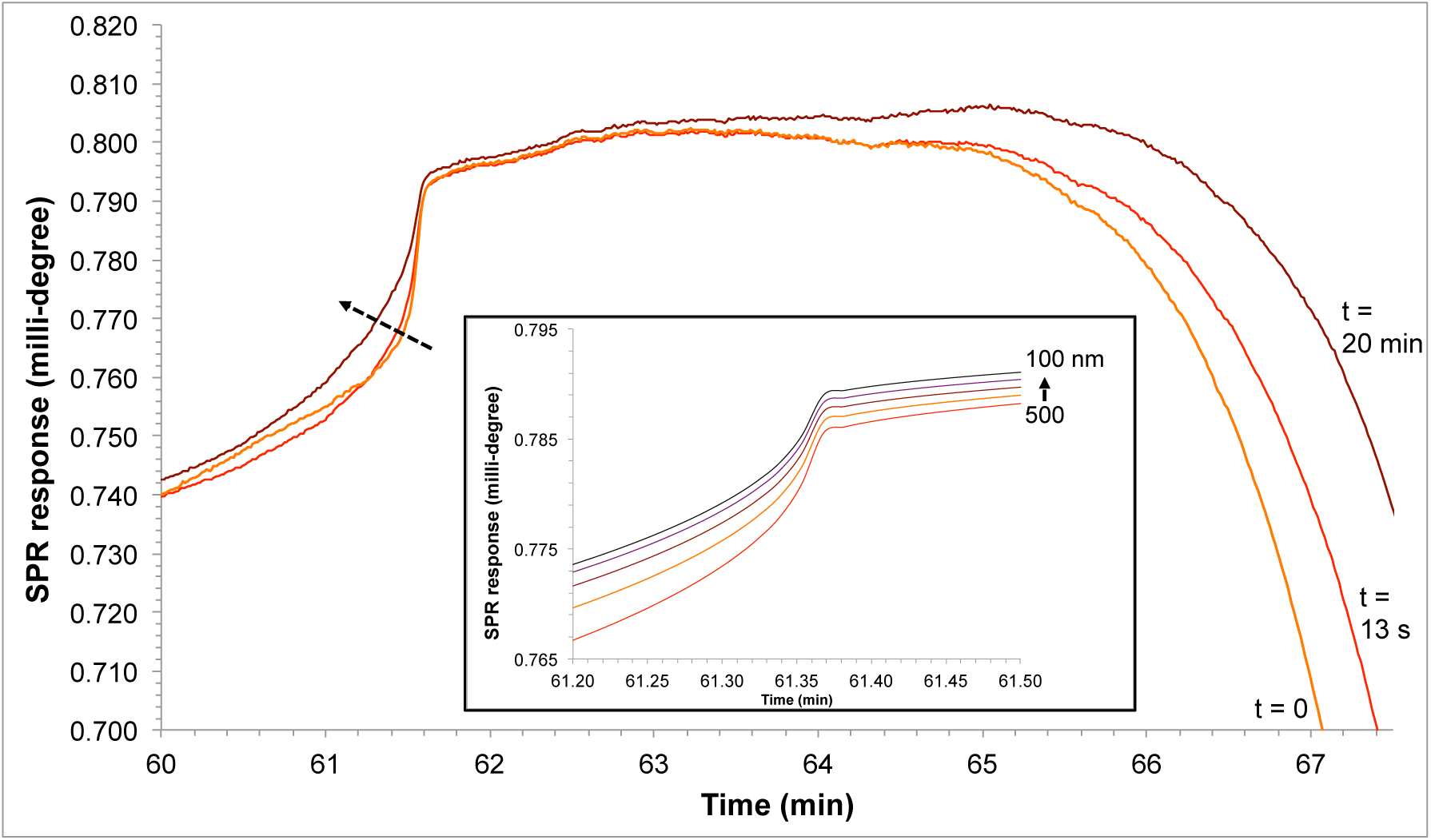
TIR detail of the full angle scans of spin-coated chitin on Au. For clarity the data derived from 670 nm is depicted, however the models were fitted for 670 and 785 nm simultaneously. In orange (t = 0) is the first time point after buffer B is introduced, in dark red (t = 20) the base line in the SPR sensorgram is stable. The dashed arrow indicates the shift of the shape of the TIR angle from thicker to a thinner layer. Inset: Layersolver model of the change of the TIR angle shape from 500 nm to 100 nm (chitin on Au).

Upon further visual inspection with SEM of the spin coated chitin layers, at first glance chitin on gold or chitin on PS before SPR measurements appear to have similar structures, though appear to be cruder on the Au-PS layer (fig. 8A and 8B). However, after one hour under normal atmospheric conditions at room temperature, chitin on gold presented large build-up structures not observed in the Au-PS layer. As seen in figure 2, this is most likely the result of the electrochemical reaction between Au and Li [51]. In order to remove the LiCl from the dried chitin layer, EDTA was added to a buffer (Buffer B). The response in the SPR sensorgram upon introduction of the liquid phase in either buffer D (without EDTA) or Buffer B (including EDTA) differed. Upon contact with buffer D, the base line in the SPR sensorgram sloped slowly down and stabilized after 15-20 minutes. In the case of the EDTA containing buffer the baseline dropped immediately and then stabilized within 20 minutes on the Au surface due to changes in the thickness (fig. 5), while the baseline is stable within seconds on the Au-PS surface. The SPR sensors with a stable baseline in buffer B were then transferred, while air-drying, within 15 minutes to the vacuum chamber of the SEM equipment (fig. 8E and 8F). The cubical structures on the Au-PS surface are clearly sodium chloride crystals [52], while it is unclear what the large macrostructures are on the Au surface at this magnification. In-between the macrostructures chitin can be seen at higher magnification (fig. 8G and 8I). On Au, chitin appears to be connected at the edge of the macrostructures (fig. 8G arrow), while chitin on the Au-PS surface appears to be more uniform.

Chitin responded to the atmospheric conditions rather rapidly, most likely due to moisture in the air, since when the Au-PS surface was directly placed in vacuum (< 30 seconds; fig. 9A and 9C) or after 5 minutes (fig. 9B and 9D), the spin-coated surfaces looked very different. At lower magnification we observed geometric patterns appear after a few minutes (fig. 9B), which were stable on the Au-PS surfaces for at least an hour (fig. 8B). At higher magnification we observed a flakey structure (fig. 9C) at first and a more uniform geometrical surface of the spin-coated surface after 5 minutes (fig. 9D).

The chitin layers on Au or Au-PS in air were similar when observed with SEM and their layer thickness was nearly the same. The differences after exposure of buffer B, at a flow of 20 μL/min in the SPR flow-cell, were evident; while the surface of Au looks damaged and not uniform, chitin on Au-PS appears smoother. The range of the chitin layer’s thickness on Au-PS was slightly lower that chitin on Au. Our layers were in the same range as earlier reported thicknesses [2].

### Differences in binding properties

The surfaces (Au spin-coated with chitin and Au coated with PS and spin-coated with chitin) did not only differ in surface properties, the chitin binding domain of *Bacillus circulans* (CDB_ChiA1_) showed different binding characteristics to chitin as well (table 2 & figure 6). CDB_ChiA1_ fused to the split-intein NpuDnaE_NΔ16_ (CBD-IN_N_) was titrated to both surfaces at increasing concentrations (figure 6). The resulting sensorgrams were then fitted to different models using the TraceDrawer software until the best fit with the lowest variation (Chi-Square) was calculated. For both experiments the best fitting model was a OneToTwo model [53]. This model indicates that the CBD-IN_N_ protein binds specifically and nonspecifically to the chitin surface.

**Table 2.** Kinetic binding properties to chitin and CBD-split intein.

| Protein | SPR Surface | KD<sub>1</sub><br>[nM] | KD<sub>2</sub><br>[nM] | B<sub>max2</sub>/B<sub>max1</sub><sup>1</sup> |
| --- | --- | --- | --- | --- |
| CBD-IN<sub>N</sub> | Au-Chitin | 76.3 | 0.0425 | 0.29 |
| CBD-IN<sub>N</sub> | Au-PS-Chitin | < 0.001<sup>4</sup> | 0.0052 | 0.08 |
| IN<sub>C</sub>-eGFP | Au-Chitin + 0.025 nM<sup>2</sup> CBD- |  |  |  |
|  | IN<sub>N</sub> | 90.3 | 0.0026 | 0.18 |
| IN<sub>C</sub>-eGFP | Au-Chitin + 0.25 nM<sup>3</sup> CBD- |  |  |  |
|  | IN<sub>N</sub> | 42.5 | < 0.001<sup>4</sup> | 0.23 |
| IN<sub>C</sub>-eGFP | Au-PS-Chitin + 0.025<sup>2</sup> nM |  |  |  |
|  | CBD-IN<sub>N</sub> | 13.8 | 0.0571 | 0.40 |
| IN<sub>C</sub>-eGFP | Au-PS-Chitin + 0.25<sup>3</sup> nM |  |  |  |
|  | CBD-IN<sub>N</sub> | 1.63 | 0.0509 | 0.48 |

(1) B<sub>max2</sub>/B<sub>max1</sub> is the ratio of non-specific binding over specific binding; (2) 0.5 µg/ml; (3) 5 µg/ml; (4) Upper limit for affinity, as precision in fits cannot be obtained due to the limited dissociation of these samples.

**Figure 6.**
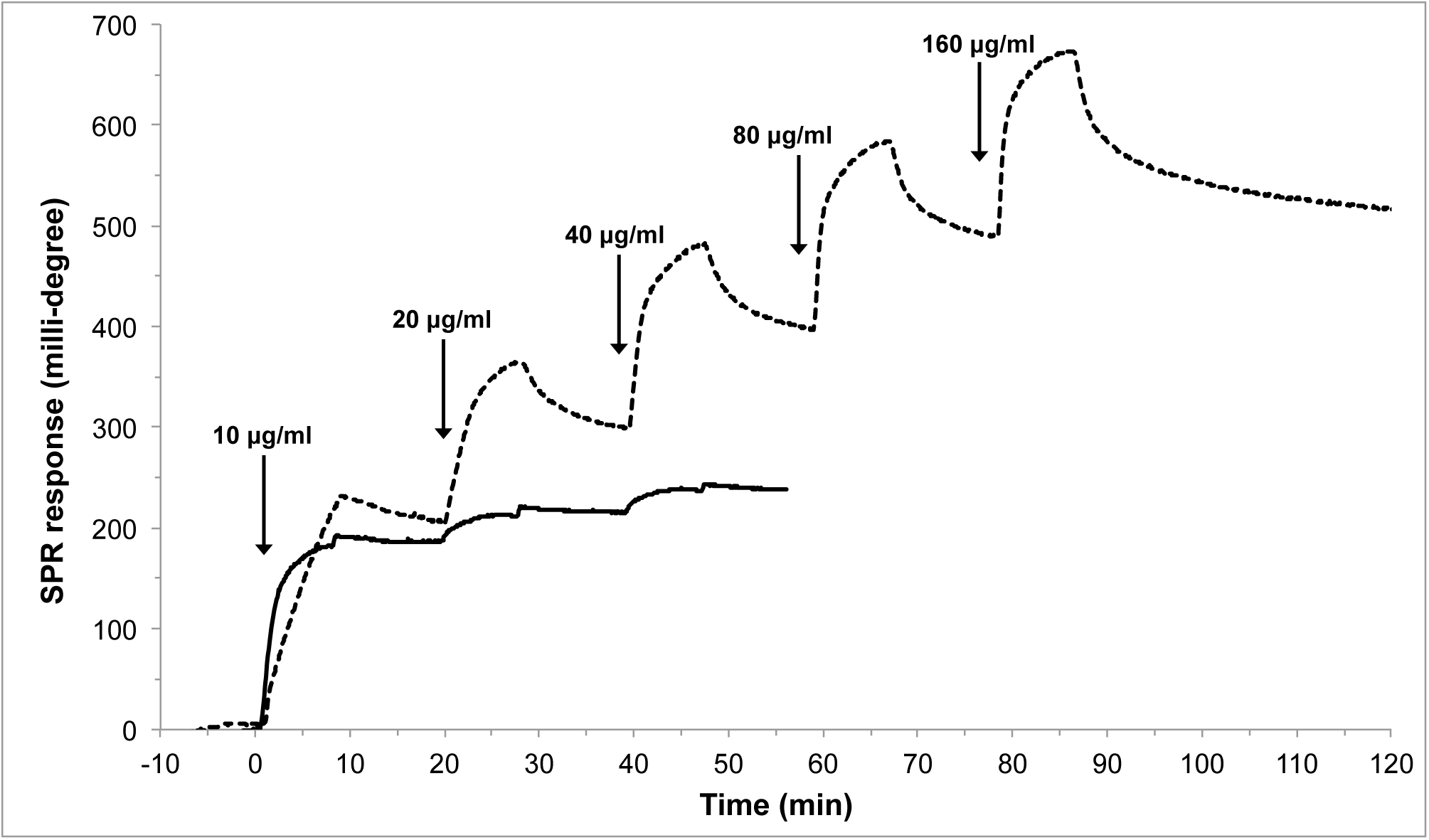
Titration of CBD-IN_N_ onto a spin-coated (SC) chitin layer on Au (dashed) or SC chitin layer on PS (solid line). Concentrations are equivalent to 490, 981, 1961, 3922, and 7854 nM.

The chitin-binding domain bound very tightly to both chitin layers and we did not observe a dissociation phase, however there was a clear difference in binding to spin-coated chitin on gold versus spin-coated chitin on a thin PS layer. Not only was the binding of the CBD to chitin on PS must stronger, the binding rates were higher, and binding was more specific (table 2 and figure 6).

The binding constants for various CBDs have been mainly determined by means of solution depletion studies and ICT (table 3). Though the theoretical differences between the solution depletion method and SPR had been thoroughly described [54, 55], to the best of our knowledge we could find only one direct comparison of both methods of a protein interaction with a polysaccharide: the study of BSA binding to cellulose. When the binding constant of BSA to cellulose was determined by isothermal titration calorimetry (ITC) and the solution depletion method at pH 7-7.5, the authors [56] found a K_d_ of 10 μM or 0.66 mg/ml by ICT and 38 μM or 2.53 mg/ml by solution depletion followed by UV spectroscopy, with a B_max_ of 0 (ICT) or 36.3 mg (solution depletion & UV) BSA/gram cellulose extrapolated from their graphs for 0% pyridinium-grafted chitin nanocrystals, while in an earlier SPR study [57], at pH 7-7.5, about 0.5 mg/cm^2^ was bound to the sensor surface. This was an equivalent to a B_max_ of about 0.018 mg BSA/gram of cellulose. The binding kinetics were not determined. The B_max_ determined with ICT of zero is close to the value determined with SPR and the difference may in the margin of error. The B_max_ determined with the solution depletion method was over 2000-fold higher than with SPR. Though the ease of the solution depletion method cannot be denied, SPR offers a more detailed analysis of the surface properties and is more sensitive. On the other hand, as pointed out by Lombardo *et al*. [56], during SPR measurements, a constant flow of BSA is supplied over immobilized cellulose, forcing the protein to adsorb to a higher extent. ITC seems to be closer in correlation with SPR than the depletion method. Therefore, when comparing the values in table 3 derived from the solution depletion method with SPR, we can expect lower K_d_ values with at least one order of magnitude. In addition, as determined by Hashimoto *et al.* [62], CBD_CHA1_ binds only to insoluble chitin, such as shrimp chitin powder. This may explain the even higher affinity (i.e. smaller K_d_) in the case of chitin spin-coated on the polystyrene surface, since the chitin layer formed a stable, uniform layer immediately upon the introduction of buffer B. Based on these observations, and an earlier one [23], we assume that the chitin structures mimic shrimp shell well enough for the study of chitin and its interaction with biomolecules, such as chitinases.

**Table 3.**
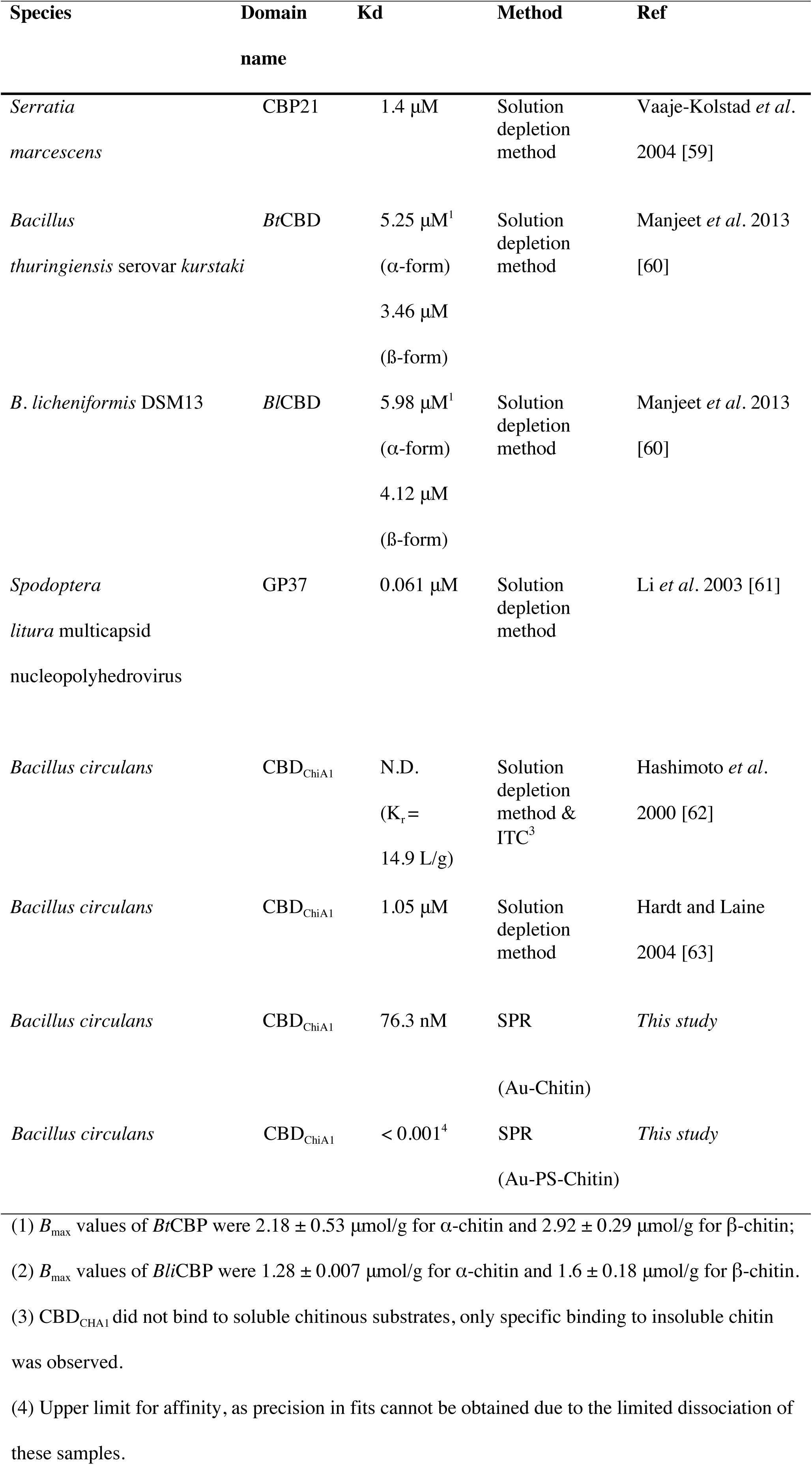
Selection of CBD and their binding characteristics with chitin.

The chitin structure had an effect on its interaction with CBD-IN_N_ and IN_C_-eGFP. Though in both the Au-chitin and the Au-PS-chitin surface the CBD-IN_N_ and IN_C_-eGFP interaction followed a capture and collapse mechanism the kinetic data fitting of our SPR experiments (Fig. 7) confirmed by the observation of the single binding phase observed with stopped-flow fluorescence [58]. The calculated binding constants were higher for the Au-chitin surface, but with a higher specific binding. On the other hand, on the Au-PS-chitin surface, the intein pair bound more tightly, but showed some non-specific binding as well. The binding constant of the split intein pair was in general higher than observed previously, except for Au-PS with a 0.25 nM CBD-IN_N_ solution titrated to the surface, before introduction of the split intein pair: 1.2 nM [58] and 1.6 nM (this study; table 2).

**Figure 7.**
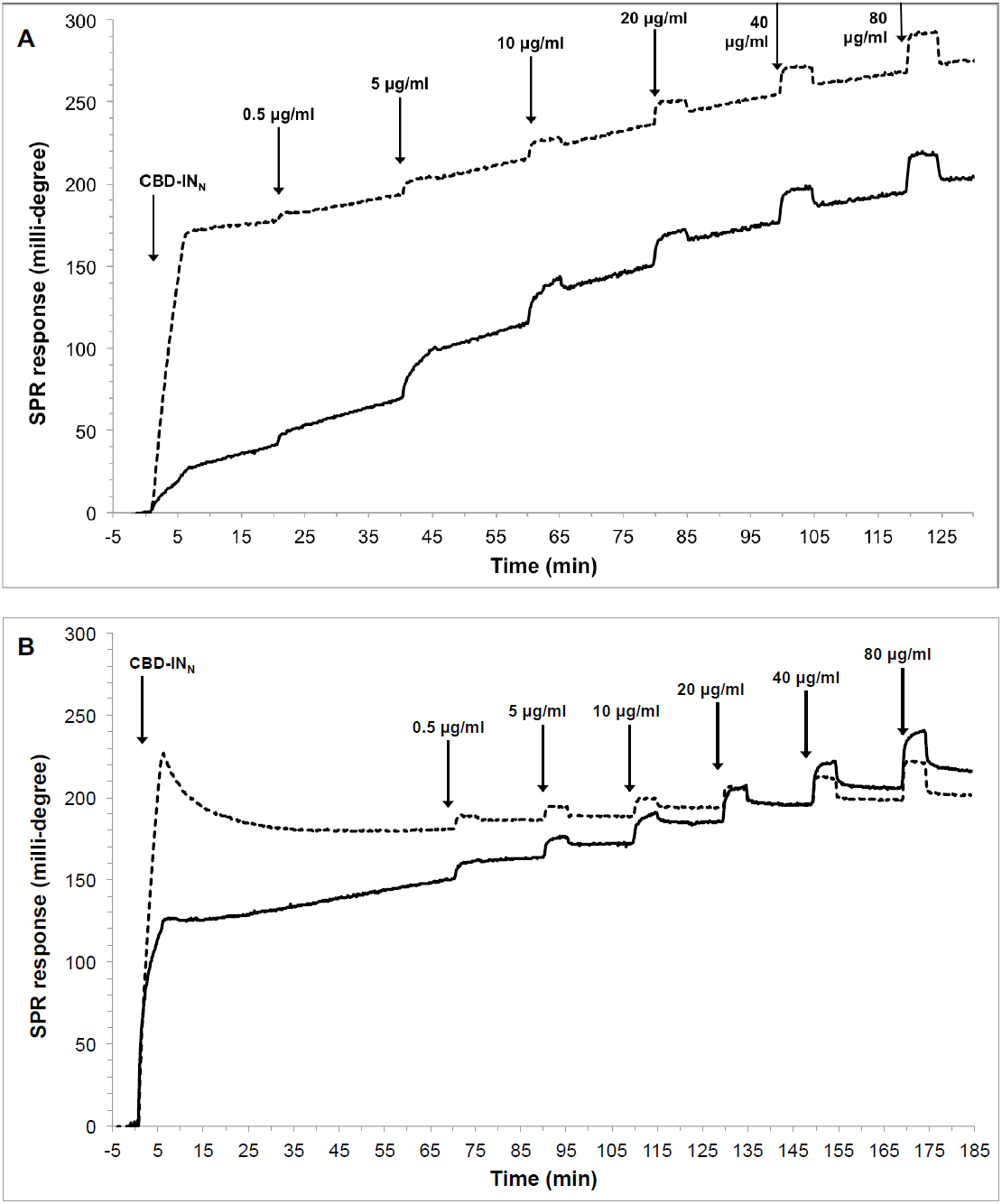
Titration of IN_c_-eGFP onto a spin-coated chitin layer on (A) Au or (B) SC chitin layer on PS after binding of 5 μg/ml CBD-IN_N_ (dashed lines) or 0.5 μg/ml CBD-IN_N_ (solid lines). Concentrations are equivalent to 17.5, 175, 350, 701, 1401, 2803 nM.

**Figure 8.**
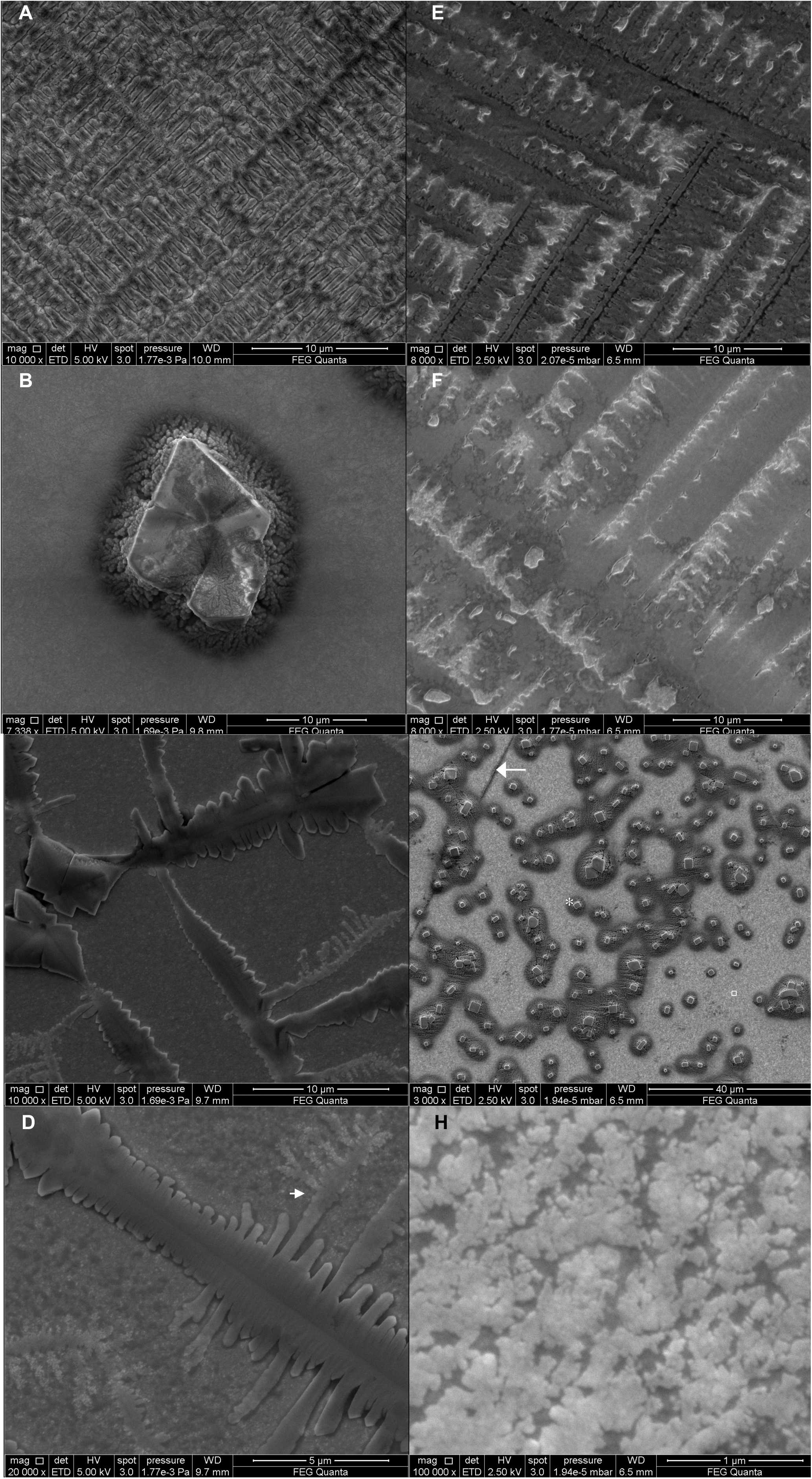
SEM images of the chitin surface (A) on Au in air (after 5 min), (B) on Au in air (after **1** hour), (C) on Au after 15 min of buffer B in the SPR flow cell (at 20 μL/min), and (D) as C, with higher resolution (arrow shows microstructure of chitin is attached to the macrostructure). (E) on Au-PS in air (5 min), (F) on PS in air (after 1 hour), (G) on Au-PS after 15 min of buffer B (at 20 μL/min); (*) cubic shapes are likely sodium chloride crystals; the arrow in F shows a line of noncoated Au, between two areas of PS coating; small white square in F the magnified area shown in G, and (H) as G, with higher resolution.

**Figure 9.**
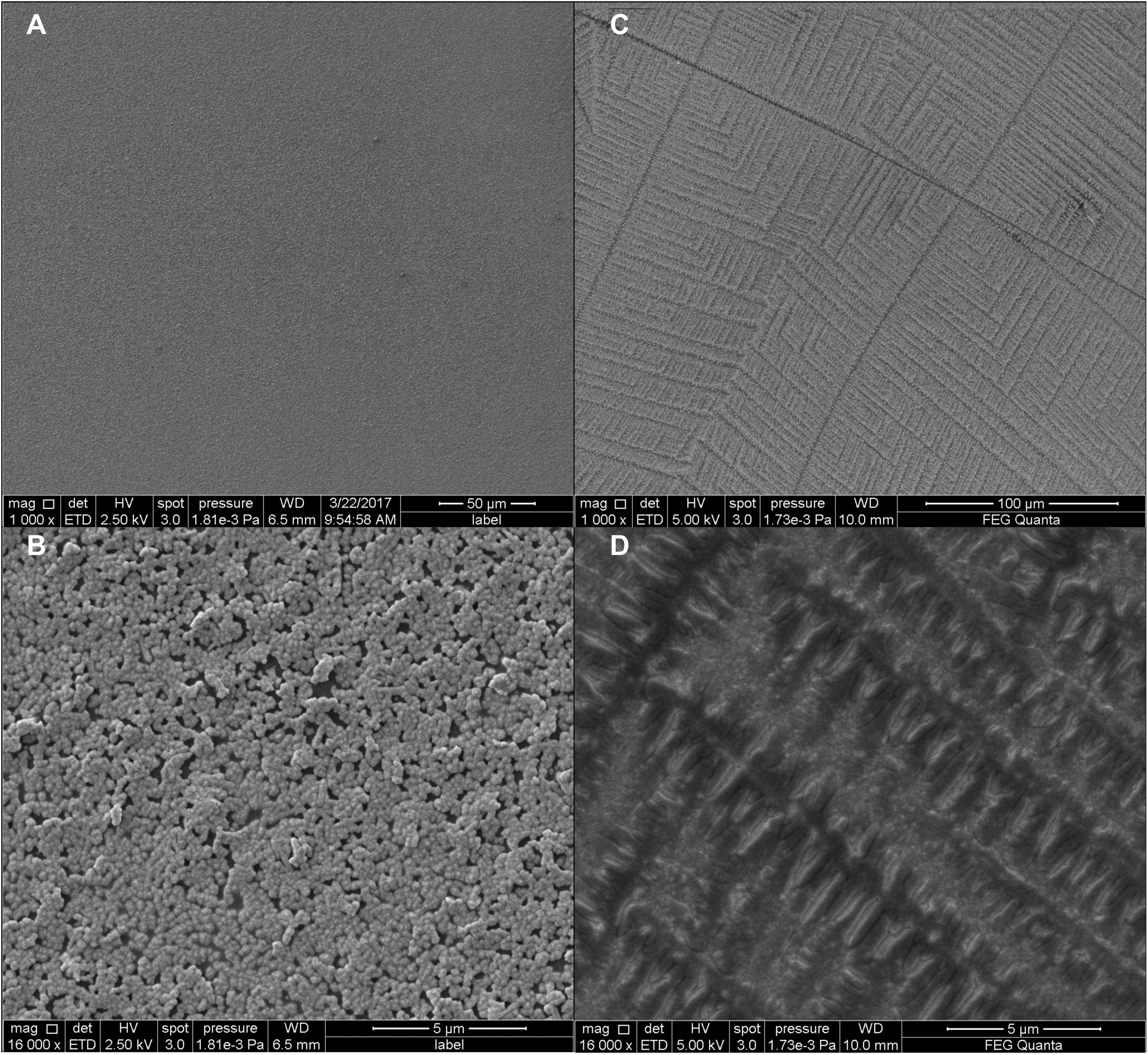
SEM images of chitin spin coated on Au-PS in air, after (A) < 30 seconds, (B) < 30 seconds at higher magnification, (C) 5 minutes, (D) 5 minutes at higher magnification.

## Conclusions

In summary, this work highlights the differences between two different sensor chips and their implication for the study of chitin binding proteins and chitin binding domain as an immobilizing agent for the study of split-inteins. The spin-coated films of chitin used with DMA/LiCl as a solvent is damaging for the gold layer, despite the use of EDTA in the SPR running buffer. Polystyrene on a gold sensor protects the gold layer from the solvent well enough for a homogeneous chitin layer to form during the spin-coating process. SPR is a suitable tool to determine the thickness of the spin-coated layers before applying buffer, during the introduction of buffer to the system, and to investigate the binding behavior of protein binding to the sensors. The utility of chitin films as biosensors is evident from the chitin binding domain binding studies. SPR is highly sensitive and new surface plasmon resonance surfaces based on copper [65] or zinc oxide [66] are an interesting avenue for future applications when combined with biomimetic approaches based on chitin metal composites of the same metals (e.g. copper [7] or zinc oxide [67]). The results we presented here are expected to enable studies of chitin layer properties and interactions with biomolecules without the use of labels in real-time with high sensitivity.

## Acknowledgments

This work was funded and supported by the Academy of Finland projects 272597, 303884, and 137053. We thank Dr. Hideo Iwai for providing us with the plasmid containing the NpuDnaE gene. We thank Dr. Ilya Belevich from the Electron Microscopy Unit, Institute of Biotechnology, of the University of Helsinki, Finland for his technical guidance with scanning electron microscopy and image analysis. We also would like to thank Erja-Liisa Piitulainen and Leena Pietilä for their kind laboratory assistance, and Regina Casteleijn-Osorno of AaltoEE and Dr. Alex Bunker of the Drug Research Program at the University of Helsinki for comments that greatly improved the manuscript.

## References

1. C.K.S. Pillai, W. Paul, C.P. Sharma, Chitin and chitosan polymers: Chemistry, solubility and fiber formation, Prog. Polym. Sci. 34 (2009) 641–678.

2. J.D. Kittle, C. Wang, C. Qian, Y. Zhang, M. Zhang, M. Roman, J.R. Morris, R.B. Moore, A.R. Esker, Ultrathin Chitin Films for Nanocomposites and Biosensors, Biomacromolecules, 13 (2012) 714–718. DOI: 10.1021/bm201631r

3. M.N.V.R. Kumar, A review of chitin and chitosan applications, React. Funct. Polym. 46 (2000), 1 – 27.

4. M. Kaya, M. Mujtaba, H. Ehrlich, A.M. Salaberria, T. Baran, C.T. Amemiya, R. Galli, L. Akyuz, I. Sargin, J. Labidi, On chemistry of gamma-chitin, Carbohydr Polym, 176, (2017), 177–186.

5. M. Wysokowski, I. Petrenko, A.L. Stelling, D. Stawski, T. Jesionowski, H. Ehrlich H, Poriferan Chitin as a Versatile Template for Extreme Biomimetics. Polymers, 7 (2015) 235–265.

6. M. Wysokowski, V.V. Bazhenov, M.V. Tsurkan, R. Galli, A.L. Stelling, H. Stocker, S. Kaiser, E. Niederschlag, G. Gartner, T. Behm, M. Ilan, A.Y. Petrenko, T. Jesionowski, H. Ehrlich, Isolation and identification of chitin in three-dimensional skeleton of Aplysina fistularis marine sponge, Int J Biol Macromol, 62 (2013), 94–100.

7. I. Petrenko, V.V. Bazhenov, R. Galli, M. Wysokowski, J. Fromont, P.J. Schupp, A.L. Stelling E. Niederschlag, H. Stoker, V.Z. Kutsova, T. Jesionowski, H. Ehrlich, Chitin of poriferan origin and the bioelectrometallurgy of copper/copper oxide. Int J Biol Macromol, 104 (2017) 1626–1632.

8. B.K. Park, M.M. Kim, Applications of Chitin and Its Derivatives in Biological Medicine. Int. J. Mol. Sci., 11(12) (2010) 5152–5164 http://doi.org/10.3390/ijms11125152

9. P. Jolles, R.A.A. Muzzarelli, Chitin and Chitinases. Jollès P, Muzzarelli RAA, editors. Birkhauser Verlag; Basel, Switzerland, 1999.

10. A. Anitha A, S. Sowmya, P.T.S. Kumar, S. Deepthi, K.P. Chennazhi, H. Ehrlich, M. Tsurkan, R. Jayakumar, Chitin and chitosan in selected biomedical applications. Progress in Polymer Science, 39 (2014), 1644–1667.

11. W. Suginta, P. Khunkaewla, A. Schulte, Electrochemical biosensor applications of polysaccharides chitin and chitosan. Chem Rev, 113 (2013) 5458–5479.

12. I. Younes, M. Rinaudo, Chitin and chitosan preparation from marine sources. Structure, properties and applications. Mar Drugs, 13 (2015) 1133–1174.

13. K.M. Rudall, W. Kenchington, The Chitin System, Biological Reviews, 48 (1973), 597–633. doi: 10.1111/j.1469-185X.1973.tb01570.x

14. J. Sakamoto, J. Sugiyama, S. Kimura, T. Imai, T. Itoh, T. Watanabe, S. Kobayashi, Artificial chitin spherulites composed of single crystalline ribbons of α-chitin via enzymatic polymerization. Macromolecules. 33 (2000) 4155–4160.

15. J. Ruiz-Herrera, V.O. Sing, W.J. van der Woude, S. Bartnicki-Garcia, Microfibril assembly by granules of chitin synthetase. Proc. Natl. Acad. Sci. USA, 72 (1975) 2706–2710.

16. S. Bartnicki-Garcia, J. Persson, H. Chanzy, An electron microscope and electron diffraction study of the effect of calcofluor and congo red on the biosynthesis of chitin *in vitro*. Arch. Biochem. Biophys. 310 (1994) 6–15.

17. W. Helbert, J. Sugiyama, High-resolution electron microscopy on cellulose II and α-chitin single crystals, Cellulose, 5 (1998) 113–122.

18. C.P. Moraru CP, V. Ostafe, Preliminary results on estimation of the degree of acetylation of chitosan by UV spectrophotometry, New Front Chem, 24 (2015), 125–133.

19. G. Vaaje-Kolstad, B. Westereng, S.J. Horn, Z. Liu, H. Zhai, M. Sørlie, V.G.H. Eijsink, Science, 330 (2010) 219–222.

20. S. Boddohi, M.J. Kipper, Engineering Nanoassemblies of Polysaccharides, Adv. Mater. 22 (2010) 2998–3016

21. I. Stepniak, M. Galinski, K. Nowacki, M. Wysokowski, P. Jakubowska, V.V. Bazheno, T. Leisegang, H. Ehrlich, T. Jesionowski, A novel chitosan/sponge chitin origin material as a membrane for supercapacitors - preparation and characterization. Rsc Advances, 6 (2016) 4007–4013.

22. E. Kim, Y. Xiong, Y. Cheng, H.C. Wu, Y. Liu, B.H. Morrow, H. Ben-Yoav, R. Ghodssi, G.W. Rubloff, J.N. Shen, W.E. Bentley, X. Shiet, G.F. Payne, Chitosan to Connect Biology to Electronics: Fabricating the Bio-Device Interface and Communicating Across This Interface. Polymers, 7 (2015) 1–46.

23. B. Focher, A. Naggi, G. Torri, A. Cosani, M. Terbojevich, Structural differences between chitin polymorphs and their precipitates from solutions—evidence from CP-MAS C-NMR, FT-IR and FT-Raman spectroscopy, Carbohydr Polym., 17 (1992) 97–102.

24. H. Ehrlich, S. Heinemann, C. Heinemann, P. Simon, V.V. Bazhenov, N.P Shapkin, R. Born, K.R. Tabachnick, T. Hanke, H. Worch, Nanostructural organization of naturally occurring composites - Part I: Silica-collagen-based biocomposites. Journal of Nanomaterials (2008) Article ID 623838, 8 pages, doi:10.1155/2008/623838.

25. H. Ehrlich H, Chitin and collagen as universal and alternative templates in biomineralization. International Geology Review, 52 (2010), 661–699.

26. E. Brunner E, P. Richthammer, H. Ehrlich, S. Paasch, P. Simon, S. Ueberlein, K.H. van Pee, Chitin-Based Organic Networks: An Integral Part of Cell Wall Biosilica in the Diatom Thalassiosira pseudonana. Angewandte Chemie-International Edition, 48 (2009), 9724–9727.

27. M. Wysokowski, K. Materna, J. Walter, I Petrenko, A.L. Stelling, V.V. Bazhenov, L. Klapiszewski, T. Szatkowski, O. Lewandowska, D. Stawski, S.L. Molodtsov, H. Maciejewski, H. Erhlich, T. Jesionowski, Solvothermal synthesis of hydrophobic chitin-polyhedral oligomeric silsesquioxane (POSS) nanocomposites. Int J Biol Macromol, 78 (2015) 224–229.

28. M. Wysokowski, T. Behm, R. Born, V.V. Bazhenov, H. Meissner, G. Richter, K. Szwarc-Rzepka, A. Makarova, D. Vyalikh, P. Schupp, T. Jesionowski, H. Erhlich. Preparation of chitin-silica composites by in vitro silicification of two-dimensional Ianthella basta demosponge chitinous scaffolds under modified Stober conditions. Mater Sci Eng C Mater Biol Appl, 33 (2013) 3935–3941.

29. v. P.P. Weimarn, Eine allgemein anwendbare Methode, Fibroin, Chitin, Kasein und ähnliche Substanzen nit Hilfe kozentrierter wässeriger Lösungen leicht löslicher und stark hydratisierter Salze in den plastischen Zustand und in den Zustand kolloider Lösung überzuführen. Kolloid-Zeitschrift, 40 (1926) 120–122.

30. v. P.P. Weimarn, Kolloide Auflösung hochmolekularer Verbindungen durch sehr leicht lösliche, stark hydratisierte Substanzen. Kolloid-Zeitschrift, 42 (1927) 134–140.

31. X. Shen, J. L. Shamshina, P. Berton, G. Gurauc and R. D. Rogers, Hydrogels based on cellulose and chitin: fabrication, properties, and applications, Green Chem., 18 (2016) 53–75.

32. C. Zhong, A. Cooper, A. Kapetanovic, Z. Fang, M. Zhang and M. Rolandi, A facile bottom-up route to self-assembled biogenic chitin nanofibers. Soft Matter, 6 (2010) 5298–5301.

33. Y. Kikkawa, H. Tokuhisa, H. Shingai, T. Hiraishi, H. Houjou, M. Kanesato, T. Imanaka, T. Tanaka, Interaction Force of Chitin-Binding Domains onto Chitin Surface, Biomacromolecules. 9 (2008) 2126–2131.

34. R.B.M. Schasfoort, A.J. Tudos, Handbook of Surface Plasmon Resonance, The Royal Society of Chemistry, Cambridge, 2008.

35. J. Sambrook, D.W. Russell, Molecular cloning. A laboratory manual, 3rd edition, Cold spring harbor laboratory press, 2001.

36. J. Panula-Perälä, J. Šiurkus, A. Vasala, R. Wilmanowski, M.G. Casteleijn, P Neubauer, Enzyme controlled glucose auto-delivery for high cell density cultivations in microplates and shake flasks, Microbial Cell Factories, (2008) 7, 31. http://doi.org/10.1186/1475-2859-7-31

37. K. Ukkonen, A. Vasala, H. Ojamo, P. Neubauer, High-yield production of biologically active recombinant protein in shake flask culture by combination of enzyme-based glucose delivery and increased oxygen transfer. Microbial Cell Factories, (2011) 10, 107. http://doi.org/10.1186/1475-2859-10-107

38. E. Gasteiger, C. Hoogland, A. Gattiker, S. Duvaud S, M.R. Wilkins, R.D. Appel, A. Bairoch, Protein Identification and Analysis Tools on the ExPASy Server, (In) John M. Walker (ed): The Proteomics Protocols Handbook, Humana Press, 2005, pp. 571–607

39. P.R. Austin. Chitin solvents and solubility parameters. J.P. Zikakis (Ed.), Chitin and chitosan and related enzymes, Academic Press, Inc., Orlando, 1984, pp. 227–237

40. S. Chong, F.B. Mersha, D.G. Comb, M.E. Scott, D. Landry, L.M. Vence, F.B. Perler, J. Benner, R.B. Kucera, C.A. Hirvonen, J.J. Pelletier, H. Paulus, M.Q. Xu, Single-column purification of free recombinant proteins using a self-cleavable affinity tag derived from a protein splicing element, Gene, 192, (1997) 271–281. DOI: 10.1016/S0378-1119(97)00105-4.

41. K. W. Bewig, W. A. Zisman, The Wetting of Gold and Platinum by Water, J. Phys. Chem., 69 (1965) 4238–4242. DOI: 10.1021/j100782a029

42. V.L. Deringer, U. Englert, R. Dronskowski, Nature, Strength, and Cooperativity of the Hydrogen-Bonding Network in α-Chitin, Biomacromolecules, 17 (2016) 996–1003. DOI: 10.1021/acs.biomac.5b01653

43. T. Kameda, M. Miyazawa, H. Ono, M. Yoshida, Hydrogen Bonding Structure and Stability of α-Chitin Studied by ^13^C Solid-State NMR. Macromol. Biosci., 5 (2005) 103–106. doi: 10.1002/mabi.200400142

44. C. C. Dupont-Gillain, Y. Adriaensen, S. Derclaye, P. G. Rouxhet, Plasma-Oxidized Polystyrene: Wetting Properties and Surface Reconstruction. Langmuir, 16 (2000) 8194–8200. DOI: 10.1021/la000326l

45. N. Granqvist, H. Liang, T. Laurila, J. Sadowski, M. Yliperttula, T. Viitala, Characterizing Ultrathin and Thick Organic Layers by Surface Plasmon Resonance Three-Wavelength and Waveguide Mode Analysis. Langmuir, 29 (2013) 8561–8571. DOI: 10.1021/la401084w

46. D.E. Azofeifa, H.J. Arguedas, W.E. Vargas, Optical properties of chitin and chitosan biopolymers with application to structural color analysis, Optical Materials, 35 (2012) 175–183.

47. K. Manabe, C. Tanaka, Y. Moriyama, M. Tenjimbayashi, C. Nakamura, Y. Tokura, T. Matsubayashi, K.H. Kyung, S. Shiratori, Chitin Nanofibers Extracted from Crab Shells in Broadband Visible Antireflection Coatings with Controlling Layer-by-Layer Deposition and the Application for Durable Antifog Surfaces, ACS App. Mat. & Inter., 8 (2016) 31951–31958.

48. J. Franklin, Z.Y. Wang, Refractive Index Matching: A General Method for Enhancing the Optical Clarity of a Hydrogel Matrix. Chem. Mat. 14 (2002) 4487–4489.

49. Labware chemical resistance table, ThermoFisher Scientific, https://tools.thermofisher.com/content/sfs/brochures/D20480.pdf (accessed 30.09.2017)

50. K. Norrman, A. Ghanbari-Siahkalia, N.B. Larsena, 6 Studies of spin-coated polymer films, Annu. Rep. Prog. Chem., Sect. C: Phys. Chem., 101 (2005) 174–201. DOI: 10.1039/B408857N.

51. A Zhiyuan Zeng, W.I. Liang, Y.H. Chu, H. Zheng, *In situ* TEM study of the Li–Au reaction in an electrochemical liquid cell. Faraday Discuss., 176 (2014) 95–107.

52. R. Tran, E. Naseri, A. Kolasnikov, X. Bai, Y. Yang, A new generation of sodium chloride porogen for tissue engineering. Biotechnol. Appl. Biochem., 58 (2011) 335–344.

53. S. Bondza, E. Foy, J. Brooks, K. Andersson, J. Robinson, P. Richalet, J. Buijs, Real-time Characterization of Antibody Binding to Receptors on Living Immune Cells. Frontiers in Immunology, (2017), 8, 455 -. DOI: 10.3389/fimmu.2017.00455

54. V. Hlady, J. Buijs, H.P. Jennissen, Methods for Studying Protein Adsorption. Methods in Enzymology, 309 (2011), 402–429.

55. E.A. Vogler, Protein Adsorption in Three Dimensions. Biomaterials, 33 (2012) 1201–1237.

56. S. Lombardo S, S. Eyley, C Schütz, H. van Gorp, S. Rosenfeldt, G. Van den Mooter, W. Thielemans, Thermodynamic Study of the Interaction of Bovine Serum Albumin and Amino Acids with Cellulose Nanocrystals. Langmuir, 33 (2017), 5473–5481.

57. H. Orelma, I. Filpponen, L.S. Johansson, J. Laine, O.J. Rojas, Modification of Cellulose Films by Adsorption of CMC and Chitosan for Controlled Attachment of Biomolecules, Biomacromolecules, 12 (2011) 4311–4318.

58. N.H. Shah, E. Eryilmaz, E.D. Cowburn, T.W. Muir, Naturally Split Inteins Assemble through a “Capture and Collapse” Mechanism. J. Am. Chem. Soc. 135 (2013) 18673–18681.

59. G. Vaaje-Kolstad, S.J. Horn, D.M.F. van Aalten, B. Synstad B, V.G.H. Eijsink, The Non-catalytic Chitin-binding Protein CBP21 from Serratia marcescens Is Essential for Chitin Degradation. J. Biol. Chem., 280 (2005) 28492.

60. K. Manjeet, P. Purushotham, C. Neeraja, A.R. Podile, Bacterial chitin binding proteins show differential substrate binding and synergy with chitinases, Microbiol. Res. 168 (2013) 461–468.

61. Z. Li, C. Li, K. Yang, L. Wang, C. Yin, Y. Gong, Y. Pang, Characterization of a chitin-binding protein GP37 of Spodoptera litura multicapsid nucleopolyhedrovirus, Virus Res. 1-2 (2003) 113–122.

62. M. Hashimoto, T. Ikegami, O. Seino, N. Ohuchi, H. Fukada, J. Sugiyama, T. Watanabe, Expression and Characterization of the Chitin-Binding Domain of Chitinase A1 from Bacillus circulans WL-12. J. of Bact. 182 (2000) 3045–3054.

63. M. Hardt, R.A. Laine, Mutation of active site residues in the chitin-binding domain ChBDChiA1 from chitinase A1 of Bacillus circulans alters substrate specificity: use of a green fluorescent protein binding assay. Arch. Biochem. & Biophys., 426 (2004) 286–297.

64. K. Somjit, K. Hara, M. Yoshimura, N. Matsuo, O. Kongpun, Y. Nozaki, Effect of shrimp chitin and shrimp chitin hydrolysate on the state of water and dehydration-induced denaturation of lizard fish myofibrillar protein. Food Sci. Technol. Res. J., 11 (2005) 106–11411.

65. P.S. Liu, H. Wang, X.M. Li, M.C. Rui, Z.H. Zeng, Localized surface plasmon resonance of Cu nanoparticles by laser ablation liquid media. Rsc Advances, 5 (2015), 79738–79745.

66. M.K. Lee, T.G. Kim, W. Kim, Y.M. Sung, Surface plasmon resonance (SPR) electron and energy transfer in noble metal-zinc oxide composite nanocrystals. Journal of Physical Chemistry C, 112 (2008) 10079–10082.

67. M. Wysokowski, M. Motylenko, H. Stocker, V.V. Bazhenov, E. Langer, A. Dobrowolska, K. Czaczyk, R. Galli, A.L. Stelling, T. Behm, L. Klapiszewski, D. Ambrożewicz, M. Nowacka, S.L. Molodtsov, B. Abendroth, D.C. Meyer, K.J. Kurzydłowski, T. Jesionowski, H. Ehrlich, An extreme biomimetic approach: hydrothermal synthesis of beta-chitin/ZnO nanostructured composites. Journal of Materials Chemistry B, 1 (2013), 6469–6476.

